# A phylogenetic codon substitution model for antibody lineages

**DOI:** 10.1101/078279

**Authors:** Kenneth B Hoehn, Gerton Lunter, Oliver G Pybus

**Affiliations:** Department of Zoology, University of Oxford, South Parks Road, Oxford, OX1 3PS, UK; Wellcome Trust Centre for Human Genetics, University of Oxford, Roosevelt Drive, Oxford, OX3 7BN, UK

## Abstract

Phylogenetic methods have shown promise in understanding the development of broadly neutralizing antibody lineages (bNAbs). However, the mutational process that generates these lineages – somatic hypermutation (SHM) – is biased by hotspot motifs, which violates important assumptions in most phylogenetic substitution models. Here, we develop a modified GY94-type substitution model that partially accounts for this context-dependency while preserving independence of sites during calculation. This model shows a substantially better fit to three well-characterized bNAb lineages than the standard GY94 model. We show through simulations that accounting for hotspot motifs can lead to reduced bias of other substitution parameters, and more accurate ancestral state reconstructions. We also demonstrate how our model can be used to test hypotheses concerning the roles of different hotspot and coldspot motifs in the evolution of B-cell lineages. Further, we explore the consequences of the idea that the number of hotspot motifs – and perhaps the mutation rate in general – is expected to decay over time in individual bNAb lineages.

## Introduction

Recent advances in sequencing technology are giving an unprecedented view into the genetic diversity of the immune system during infection, especially in the context of chronic infections caused by viruses. Broadly neutralizing antibody (bNAb) lineages, which produce B cell receptors (BCRs) capable of binding a wide range of viral epitopes, are of particular interest (Haynes et al. 2012). Within such lineages, all B cells descend from a shared common ancestor and are capable of rapid sequence evolution through the processes of somatic hypermutation (SHM) and clonal selection. For chronically infecting viruses such as HIV-1, this co-evolutionary process may continue for years (Wu et al. 2015). Because immunoglobulin gene sequences from bNAb lineages undergo rapid molecular evolution, selection and diversification, they would appear to be suitable for evolutionary and phylogenetic analysis, and these methods have already been applied to various immunological questions such as inferring the ancestral sequences of bNAb lineages (Sok et al. 2013; Hoehn et al. 2016). Intermediate ancestors of B cell lineages are of particular interest because they may act as targets for stimulation by vaccines (Haynes et al. 2012).

However, the biology of mutation and selection during somatic hypermutation is different from that which occurs in the germline, and therefore it is unlikely that standard phylogenetic techniques will be directly applicable to studying bNAb lineages without suffering some bias and error. One of the most important assumptions of likelihood-based phylogenetics is that evolutionary changes at different nucleotide or codon sites are statistically independent. Without this assumption, likelihood calculations become computationally impractical as the length and number of sequences increases (Felsenstein 1981). Unfortunately, in contrast to germline mutations, somatic hypermutation of BCR sequences is driven by a collection of enzymes that cause some sequence motifs (between two and seven base pairs) to mutate at a higher rate than others (Smith et al. 1996; Teng and Papavasiliou 2007; Elhanati et al. 2015). This context sensitivity clearly violates the assumption of independent evolution among sites. Furthermore, because hotspot motifs are, by definition, more mutable than non-hotspot motifs, their frequency within a B-cell lineage may decrease over time as they are replaced with more stable motifs (Sheng et al. 2016). These changes will not be passed on to subsequent generations through the germline because the mutational process is somatic. This effect may have a number of consequences for molecular evolutionary inference, for example it may render inappropriate the common practice of estimating equilibrium frequencies from the sequences themselves. At present it is unknown how the violation of these assumptions will affect phylogenetic inference of BCR sequences in practice, and the problem of ameliorating such effects remains an open issue.

This work has two main aims. The first is to analyse BCR evolution in three previously published and long-lived bNAb lineages in HIV-1 infected patients. This analysis confirms the prediction of a decay of certain hotspot motifs through time. Our second aim is to develop and introduce a new substitution model that can partially account for this effect. The model is a modification of the GY94 (Goldman and Yang 1994) codon substitution model. Although only an approximation, our new model can detect and quantify the effect of somatic hypermutation on BCR sequences whist preserving the assumption of independence among codon sites in order to maintain computational feasibility. This model shows a significantly better fit than the standard GY94 model to all three bNAb lineages from HIV-1 patients. Through simulations, we validate the effectiveness of the model, and show its ability to reduce bias in the estimation of other evolutionary parameters such as tree length. Further, we use this model as a framework for testing hypotheses of hotspot motif symmetry and hierarchy of mutability, and we explore its potential applications such as improved ancestral state reconstruction.

## Materials and Methods

### Multiple Sequence Alignment

Heavy chain sequences from the three bNAb lineages presented in (Wu et al. 2015) were downloaded from GenBank (http://www.ncbi.nlm.nih.gov/genbank/). The lineage of greatest duration was VRC01, which was sampled over 15 years (Wu et al. 2015), followed by CAP256-VRC26 (hereafter VRC26), which was sampled over four years (Doria-Rose et al. 2014), and CH103, which was sampled over three years (Liao et al. 2013). Sequences from each bNAb lineage were translated into amino acids, aligned to their putative germline V gene segment using IgBlast (Ye et al. 2013), and then re-translated back into codons. Putative germline segment assignments (V4-59*01 for CH103, V3-30*18 for VRC26, and V1-2*01 for VRC01) were obtained from bNAber (Eroshkin et al. 2013) and sequences were obtained from the IMGT V-Quest human reference set (Lefranc and Lefranc 2001). Because of considerable uncertainty in D and J germline assignments for each lineage, only the V segment was used. Insertions relative to the germline sequence were removed, so that all sequences within each lineage were aligned to the same germline sequence. Removing these insertions brought together two nucleotides that are not actually adjacent, creating false motifs. To prevent this, the 3’ nucleotide of the region joined together from the removal of the insertion was converted to an N. To keep results consistent among lineages, only nucleotide positions from the beginning of the first framework region (FWR1) to the end of FWR3 were used. Sampling dates of each sequence were extracted from the sequence ID tags provided on GenBank. Eleven sequences were excluded from CH103 because this information was not available.

### Hotspot decay in bNAb lineages

The “hotspot frequency” of each sequence was defined as the number of times a particular hotspot motif was observed, divided by the number of possible hotspot locations (sequence length - motif length + 1) in that sequence, and was calculated for two trimer (WR<u>C</u>/<u>G</u>YW) and two dimer (W<u>A</u>/<u>T</u>W) motifs separately (Yaari et al. 2013), where W = A or T, Y = A or G, and R = T or C, as per the IUPAC nucleotide ambiguity codes. Hence an example of a trimer motif might be AT<u>C</u>, and its reverse complement <u>G</u>AT. The underlined base in each of these motifs experiences increased AID-mediated mutability. Trimers and dimers with non-ACGT characters were excluded from the calculation of hotspot frequency.

Changes in hotspot frequency values through time were analysed using linear regression and correlation. Because the date of infection was not known for VRC01, germline IGHV sequences were not included in these calculations. Importantly, because the sequences within each B-cell lineage are phylogenetically related, they are partially correlated due to shared common ancestry and are not independent data points, hence *p*-values from standard correlation and regressions tests are not reliable. However, the regression is still an unbiased measure of trends in sequence change over time (see Drummond et al. 2003 for discussion). Regressions of hotspot frequency through time are shown in **Figure 1**.

**Figure 1:**
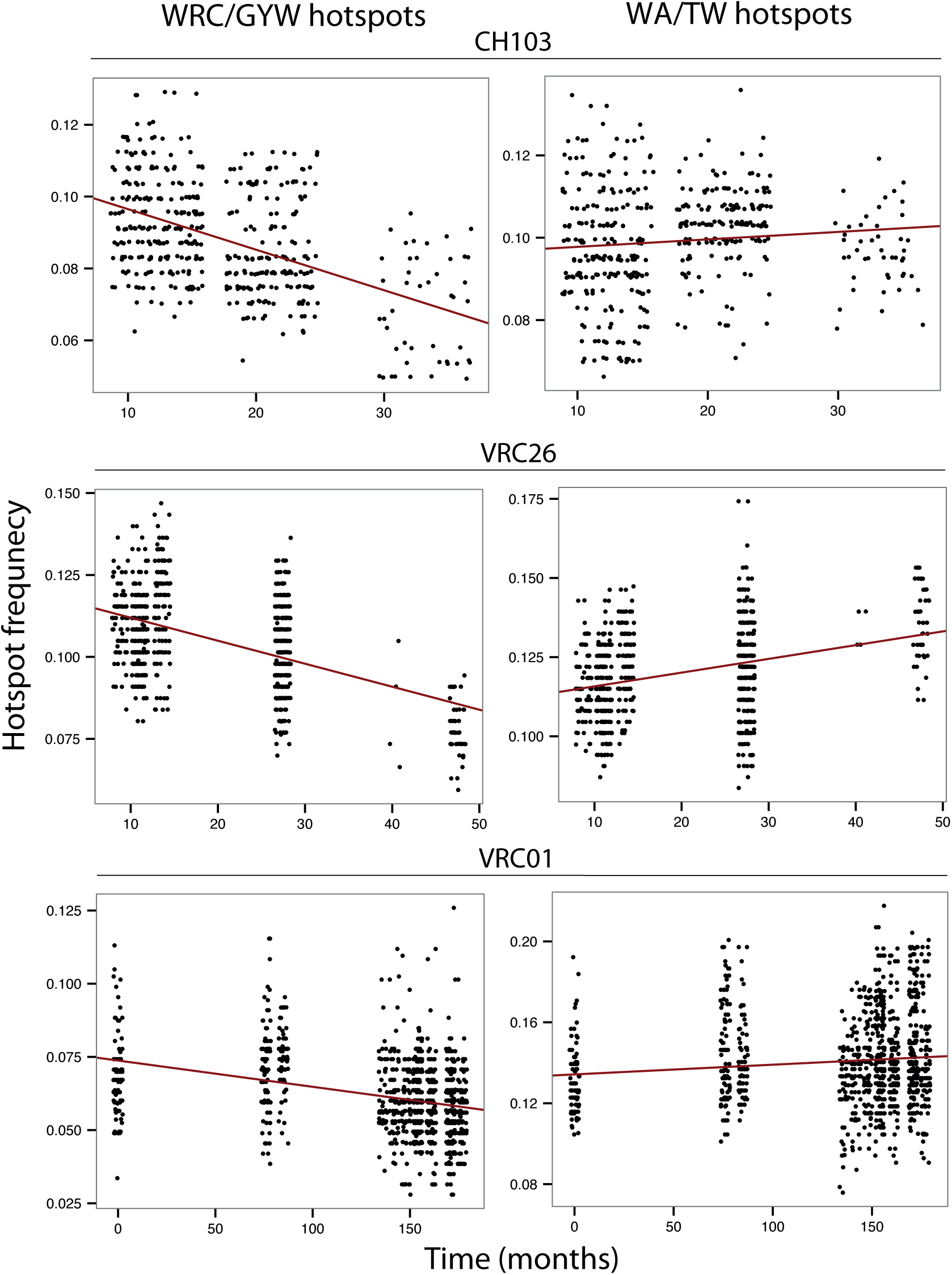
Decrease in frequency of trimer and dimer hotspot motifs in three bNAb lineages. Red line shown is least square regression.

In the absence of a suitable hypothesis test based on regression, we developed a simulation-based approach to test for significant associations between hotspot frequency and time in bNAb lineages. The null model for this test is a substitution model (GY94) that does *not* explicitly model the decay of hotspot motifs. The GY94 model is used to estimate a maximum likelihood phylogenetic tree. Multiple data sets were simulated under this null model, using the same sample sizes and sampling times as the three empirical bNAb data sets. The significance of the difference between the null model and the observed data is calculated as the proportion of simulated datasets with a greater negative correlation between hotspot frequency and time than in the observed data set. Results for these tests are shown in **Table 1**.

**Table 1:** THotspot motif decay in three bNAb lineages.

| B-cell lineage | Trimer motifs: WRC/GYW |  |  | Dimer motifs: WA/TW |  |  |
| --- | --- | --- | --- | --- | --- | --- |
|  | Observed correlation | Observed/simulated | P value | Observed correlation | Observed/simulated | P value |
| CH103 | -0.48 | -11.33 | 0.00 | 0.09 | 0.46 | 0.29 |
| VRC26 | -0.50 | -11.77 | 0.00 | 0.33 | 0.84 | 0.30 |
| VRC01 | -0.33 | 5.50 | 0.02 | 0.11 | 0.70 | 0.39 |

The “Observed correlation” column shows the correlation between hotspot frequency and time. The next column shows how these values compare to the mean of the same values from 100 simulations under the null model. The third column shows the p value – the proportion of simulated data sets that had a lower correlation than observed data sets.

Maximum likelihood phylogenetic trees and substitution model parameters for each of the three bNAb lineages were estimated using the GY94 model and empirical codon frequencies, as implemented in codonPhyML (Gil et al. 2013). Trees were re-rooted so that the germline sequence is placed as an outgroup with a branch length of zero, effectively making it the ancestor of the lineage. For each bNAb lineage, we then simulated 100 sequence data sets down the corresponding ML tree using the GY94 model, starting with the corresponding germline sequence at the root and using the fitted substitution model parameters. Simulations were performed using the program EVOLVER, which is part of the PAML package (Yang 2007).

To ascertain whether the observed effects were general, or specific to known hotspot motifs, we repeated the above regression and simulation approach for non-hotspot motifs. To do this, we simply randomly assigned non-hotspot nucleotide motifs as “hotspots” whilst keeping the number of trimer and dimer hotspots the same (eight and three, respectively). This analysis was then repeated for 100 such random allocations.

### A codon substitution model for antibody lineages

In order to represent the molecular evolution of long-lived B cell lineages more accurately, we develop here a new substitution model that models the effects of motif-specific mutation across BCR sequences. This model, named the HLP16 model, is a modification of the GY94 substitution model (more specifically, it is a modification of the M0 model, because ω is kept constant among sites and lineages; Yang et al. 2000). Specifically, we add to the GY94 model an additional parameter, *h*^*a*^, which represents the change in relative substitution rate of a hotspot/coldspot mutation in motif *a*. Explicitly modelling the full context dependence of hotspot motifs would make likelihood calculations computationally infeasible. Instead, we weight *h*^*a*^ by 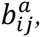 which is the probability that the mutation from codon *i* to codon *j* was a hotspot mutation in motif *a*, averaged across all possible combinations of codons on the 5’ and 3’ flanks of the target codon. This is a mean field approximation (i.e. the expected effect is averaged across all possible scenarios) and is similar to the singlet-doublet-triplet model of Whelan and Goldman (2004). A “hotspot mutation” is defined as a mutation occurring within the underlined base of the specified motif (e.g. the trimer motif and its reverse complement WR<u>C</u>/<u>G</u>YW; nucleotides represented using the IUPAC coding scheme). Because we did not find a significant decay of dimer hotspot motifs through time (**see Figure 1 and Table 1**), our model only includes trimer hotspots. However, dimers or other motifs could easily be added with additional values of *h*^*a*^ and 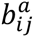 for each new motif.

In the HLP16 model, each entry *q*_*ij*_ in the transition rate matrix **Q** is parameterised by:

π_j_ = Baseline frequency of codon j

k = Transition/transversion mutation relative rate ratio

ω = Nonsynonymous to synonymous mutation relative rate ratio

a = Motif in which mutation rate is modified at underlined base. Here, *a* ∈ {WR<u>C</u>, <u>G</u>YW, W<u>A</u>, <u>T</u>W, SR<u>C</u>, <u>G</u>RS}, but in principle any other motif ≤ 4nt long could be used.

h^a^ = Change in mutability due to mutation in motif *a*; h^a^ ≥ -1.

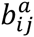 = Probability that mutation from *i* to *j* involves the underlined base in motif *a* and the transition matrix **Q** itself is defined by

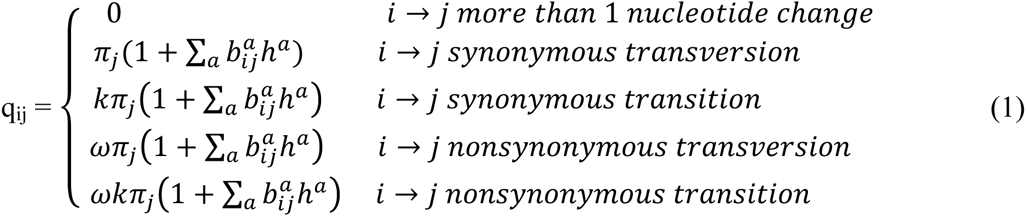

The values of 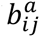 are calculated by marginalizing over all possible 5’ and 3’ flanking sense codons as follows:

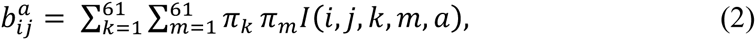

where *I* is the indicator function:

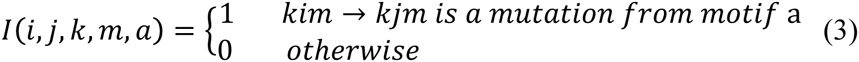

This model, though an approximation, has several useful properties. Most importantly, because codon changes are modelled as occurring independently of each other, the phylogenetic likelihood can still be calculated using Felsenstein’s pruning algorithm, which greatly reduces computational time (Felsenstein 1981). The model also has the intuitive property that, if no hotspot motif is specified, then all *h*^*a*^ = 0 and the model simplifies to the GY94 model. Thus the M0 submodel of the GY94 model is a special case of the HLP16 model.

In contrast to most substitution models, the relative substitution rate parameters in the **Q** matrix of the HLP16 model is not necessarily time-reversible, i.e. it does not necessarily satisfy the detailed balance condition π_*i*_*q*_*ij*_ = π_*i*_*q*_*ji*_. Time reversibility is useful because it means that likelihood calculations can be undertaken on an unrooted tree, which can then be rooted on any branch. In the case of B cell lineage evolution, it is necessary to root the lineage phylogeny at the germline sequence during parameter estimation. This property is also known as the “pulley principle”, which only holds for reversible models, and helps to speed up search algorithms for maximum likelihood trees (Boussau and Gouy 2006). In our implementation, likelihood calculations during branch length optimization are sped up by starting the pruning algorithm calculations at the lower (more ancestral) node of the branch being optimized, then updating the partial likelihoods on all nodes between the branch being optimized and the root node.

While in standard GY94-type models the vector **π** represents the equilibrium frequencies of codons, this is not the case for the HLP16 model. This can be checked by direct calculation of the total flux in and out of a codon *j*; in general 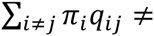 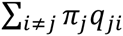 for HLP16 because the matrix 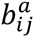 is generally not symmetric in *i* and *j*. Although equilibrium frequencies do exist (and can be calculated numerically), we are in fact interested in the model’s non-equilibrium behaviour, since the ancestral sequence is likely to be far from equilibrium, and observed codons are unlikely to have reached their equilibrium frequencies. As a result, the best-fit values of **π** may even change according to the time at which a B-cell lineage is sampled. Thus the values of **π** in our model are more appropriately interpreted as best-fit constant codon frequencies given the data and other model parameters, and should not be directly interpreted as equilibrium frequencies. More specifically, we use the CF3X4 model (Kosakovsky Pond et al. 2010) to find the best-fitting codon frequencies. In this model, the frequencies of A, C, G, and T at each of the three codon positions are estimated through ML as twelve additional parameters.

Within this framework, a hierarchical network of hotspot models can be specified by fixing certain values of *h*^*a*^ to zero and by setting some values of *h*^*a*^ to be equal. For instance, a symmetric WR<u>C</u>/<u>G</u>YW model is specified by setting *h^WR<u>C</u>^* = _*h^<u>G</u>YW^* and by setting all other values of *h*^*a*^ to zero, leaving just one parameter (*h^WR<u>C</u>^*) to be estimated using maximum likelihood. Pairs of models that are nested (e.g. strand symmetric vs. asymmetric motifs) can be formally compared using likelihood ratio tests; non-nested models may be compared using the Akaike information criterion (AIC).

We implement this model in IgPhyML, a program modified from the source code of codonPhyML (Gil et al. 2013). IgPhyML implements the rate matrix in equation 1 estimates the parameters *h*^*a*^ using maximum likelihood, together with the other model parameters. Specifically, we optimize ω, k, π_j,_ and the vector of phylogeny branch lengths. Performing all likelihood calculations from the root node slows computation substantially, therefore in this work we applied the HLP16 model to a fixed tree topology, and we deliberately leave the problem of co-estimating topology for future work. For each data set, the tree topology used was that inferred using the standard M0 version of the GY94 model in codonPhyML, which was subsequently re-rooted in order to place the germline sequence at the universal common ancestor.

Because the M0 version of the GY94 model is a special case of the HLP16 model (i.e. when all *h* parameters = 0) the two models are nested and can be compared using a likelihood ratio test. Let *L*_*max*_(*HLP16*) and *L*_*max*_(*M0*) be the maximum likelihoods obtained under the HLP16 and M0 models, respectively. The likelihood ratio statistic 2 log[*L*_*max*_(HLP16) / *L*_*max*_(M0)] is then approximately chi-squared distributed with degrees of freedom equal to the number of additional *h* parameters (Huelsenbeck and Rannala 1997). For each bNAb dataset, we calculate *L*_*max*_(*HLP16*) by co-optimising *h* and other model parameters, whereas *L*_*max*_(*M0*) is calculated by constraining all *h^a^=*0 whilst optimising the other model parameters.

### Effectiveness of the mean field approximation

We evaluated and validated our implementation of the HLP16 model by simulating data sets under different values of *h* and testing how accurately model parameters were inferred. For brevity, we considered only symmetric WR<u>C</u>/<u>G</u>YW hotspot motifs in this analysis (*h^WR<u>C</u>^=h^<u>G</u>YW^*; hereafter in this section hereafter referred to as *h*). Because the HLP16 model is a mean field approximation it will not fully account for the context dependency of somatic hypermutation. To measure the degree of this effect, we generated simulated datasets using a modified version of HLP16 that *does* fully account for the context dependence of adjacent codon sites. In a forward simulation procedure, the 3’ and 5’ flanking codons of each site are known. This allowed us to create a **B** matrix for each site in each sequence with *b*_*ij*_ equal to either 1 or 0 depending on whether or not the substitution was a hotspot mutation in a WR<u>C</u>/<u>G</u>YW motif. The process begins at the root sequence, calculates a separate **B** and **Q** matrix at each site in the sequence, simulates two descendant sequences, then repeats for descendant nodes down the tree until all tips are filled. More specifically:

1. We randomly subsampled each bNAb lineage to 99 sequences, plus the single germline sequence at the root. Subsampling was necessary to make the large number of replicates computationally feasible.
2. We estimated a maximum likelihood phylogeny for each subsampled bNAb lineage data set using the standard GY94 model. During estimation we optimised ω, k, π_j_, branch lengths and the tree topology. The resulting ML tree was re-rooted at the germline sequence with a branch length of zero.
3. For each value of *h* investigated (0, 1, 2, and 4), we simulated 20 alignments along each of these trees using the procedure outlined above. Simulations were undertaken using the estimated values of ω, k and π_j_, obtained in step (2) for the corresponding bNAb lineage data set. Starting (root) sequences were generated randomly from codon frequencies.
4. For each of the replicates defined in step (3), we performed three different ML calculations: (i) *h* was optimised using ML (with *ĥ* as the MLE estimate of *h*), (ii) *h* was fixed to zero and (iii) *h* was fixed to the true value used in simulation. These three scenarios enable us to test type 1 and type 2 error rates, by determining whether *ĥ* was significantly different to *h* or to zero, respectively. Statistical significance was determined using the chi-squared approximation to the likelihood ratio statistic, as described above. In all calculations, the tree topology was fixed to that inferred in step (2).
5. For each data set and for each set of simulations under a particular value of *h*, we estimated *ĥ* and then calculated the properties of this estimator as follows:
  i. Bias in estimation: (Mean [*ĥ*] − *h*)
  ii. Variance in estimation: Variance [*ĥ*]
  iii. Type 1 error rate: The proportion of simulated data sets in which *h* was outside of the 95% confidence interval for *ĥ*. Type 2 error rate: The proportion of simulated data sets in which *h* > 0, but failed to reject the null hypothesis (*h* = 0).

To test how our implementation performs on simulations in which HLP16 is the true model, we also repeated the above simulation analysis using the standard HLP16 model (the results of which are detailed in **Supplemental File 2**).

Biased mutation during somatic hypermutation has been shown to give false signatures of natural selection using approaches that compare the expected number of replacement and silent mutations (Dunn-Walters and Spencer 1998). We hypothesised that the HLP16 model might partially reduce this bias. To test this, and to explore whether the HLP16 model improved estimation of other evolutionary parameters, we compared the percentage error under the HLP16 and GY94 models of estimates of (i) ω, (ii) k, (iii) tree length (sum of all branch lengths) and (iv) the ratio of internal to external branch lengths. These results are provided in **Figure 2** and **Supplemental Figure 4**.

**Figure 2:**
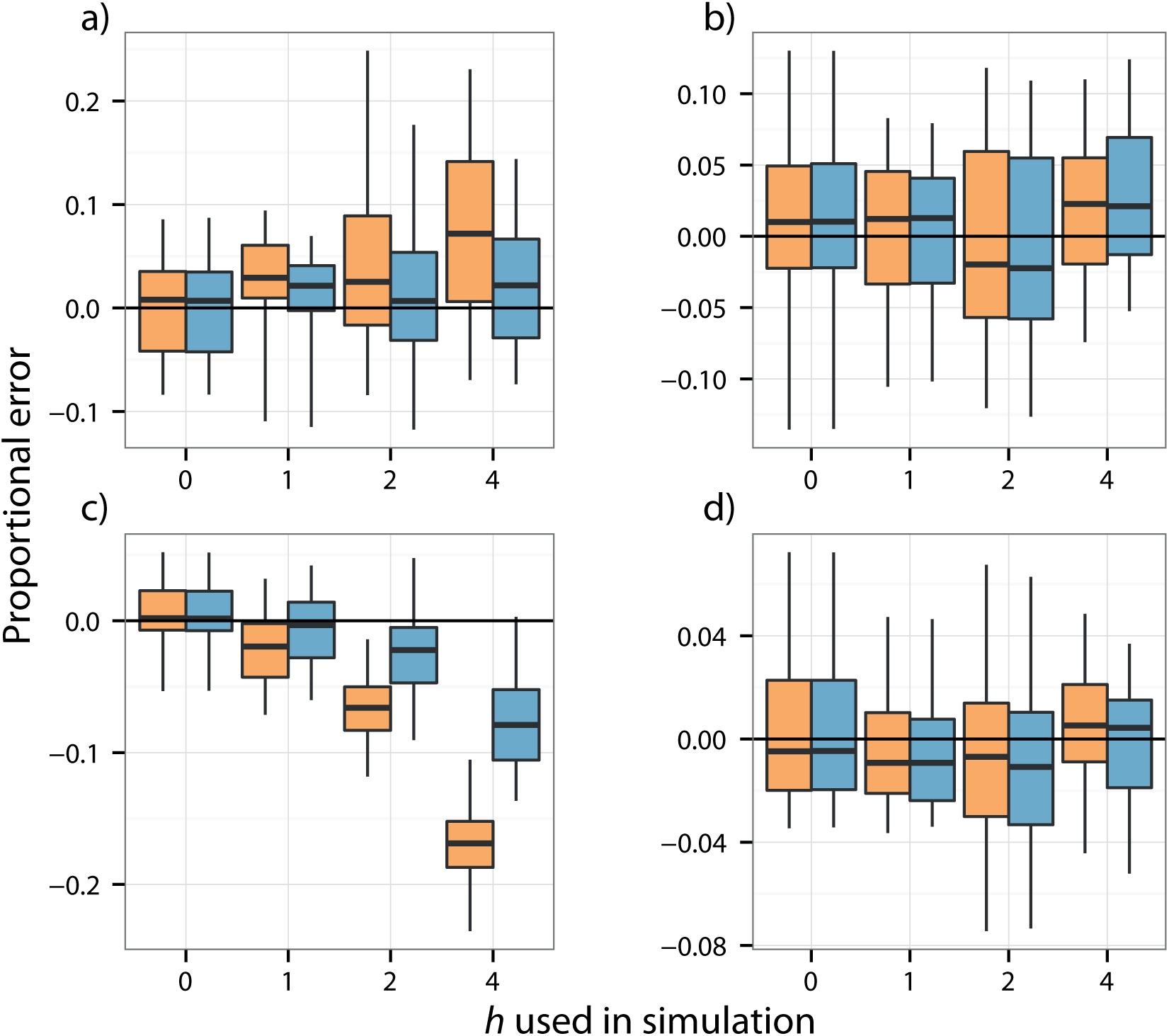
Proportional error in parameter estimation compared to true values for the VRC01 B cell lineage fully context dependent simulations. Values of ω, *k*, tree length, and ratio of internal to external branch lengths are shown in panels a), b), c), and d), respectively. Estimates obtained under the GY94 are in orange (*h=*0) and estimates obtained under the HLP16 model are in blue (*h* estimated using maximum likelihood). The edges and centres of boxplots show the 1^st^, 2^nd^, and 3^rd^ quartiles, while the whiskers show range. Similar results for B cell lineages CH103 and VRC26 are shown in **Supplemental File 4**.

The fact that bNAb lineages are clearly not in equilibrium when they are sampled (**Figure 1**) has interesting implications for the use of Markov substitution models. Typically, it is assumed that nucleotide or codon frequencies are at equilibrium at the time of sampling, and empirical codon frequencies are often used as estimates of equilibrium frequencies. In the case of long-lived B cell lineages, however, sampled sequences are almost certainly not in equilibrium, making empirical codon frequencies inaccurate approximations for equilibrium frequencies. Because changes from SHM are not inherited through the germline, each BCR lineage is expected to begin out of sequence equilibrium, potentially converging to its equilibrium distribution as it evolves. For this reason, it is necessary to optimize equilibrium codon frequencies using ML rather than using empirical codon frequencies. To test how this might affect estimation of *h*, we repeated the simulation procedure above using empirical equilibrium frequencies from each data set. These results are included in **Supplemental File 3**.

### Hotspot model selection

By placing different constraints on the six *h*^*a*^ parameters, we tested ten different hotspot models on the three bNAb lineages CH103, VRC26, and VRC01. The specific constraints used to define each hotspot model, and the results of model testing are shown in **Table 4**. Full results from each model fit are shown in **Supplemental File 5**.

Further, to ensure that the effects we observe are particular to the hotspot and coldspot motifs under investigation, we compared estimated *h* values for defined hotspot motifs to those obtained from all other possible trimer motifs with similar characteristics. Specifically, we generated all possible motifs and their reverse complements that (i) were 3nt in length, (ii) contained two IUPAC letters standing for two possible nucleotides (R, Y, S, W, K, and M), and (iii) subsequently contained an unambiguous nucleotide (i.e. A, C, G, or T). We then fitted the HLP16 model using these each of these 144 motifs individually and compared how estimated *h* values for these motifs compared the values for WR<u>C</u>/<u>G</u>YW and SY<u>C</u>/<u>G</u>RS. We repeated this process for dimer motifs, but with the constraints that motifs (i) were 2nt in length, (ii) contained one IUPAC letter standing for two possible nucleotides and (iii) subsequently contained an unambiguous nucleotide. We fitted the HLP16 model to the same data using these 24 dimer motifs and compared them to the results from W<u>A</u>/<u>T</u>W motifs. Results from this analysis are shown in **Supplemental File 6**.

### Effects on ancestral state reconstruction

One of the key applications of molecular phylogenetics to BCR sequence data is the reconstruction of ancestral sequences within a B-cell lineage (Kepler 2013). Ancestral state reconstruction is an implicit part of the phylogenetic likelihood calculation when nucleotide or codon substitution models are used. For each simulation replicate, and for each of the three likelihood calculations described in step (3) above, we computed the most likely codon at each codon position at each internal node in the tree. These ancestral sequences were then used to compare the accuracy of reconstructions under the HLP16 model with those obtained using the GY94-type model. In each simulation replicate, accuracy of ancestral sequence reconstruction was measured by calculating the mean number of pairwise nucleotide or amino acid differences between the predicted and true sequences at each node. We repeated this ancestral state reconstruction procedure on each bNAb lineage with its best-fit model. These are shown in **Supplemental File 7** and **8**, respectively.

The HLP16 model is implemented in IgPhyML, which is available to download through: https://github.com/kbhoehn/IgPhyML. Code and sequence alignments for simulation and ancestral sequence reconstruction analyses are included in the file **Supplemental_Code.zip**.

## Results

### Decay of hotspot motifs in bNAb lineages

All three bNAb lineages showed a negative correlation between trimer hotspot content and time. However, no such decline was seen in dimer motifs (**Table 1, Figure 1**). To test whether the observed patterns of hotspot decay were significantly different from those expected under a standard reversible codon substitution model that does not explicitly account for hypermutation at hotspot motifs, we implemented a significance test that compares the correlation between hotspot motif frequency and time in simulated data sets generated under the null phylogenetic model. All three B cell lineages showed a significantly greater negative correlation between trimer hotspot content and time than expected under the null model (**Table 1**). In all cases, the frequency of dimer motifs showed no significant change through time. Furthermore, we repeated these analyses with randomly chosen non-hotspot motifs taking the place of the real, known hotspot motifs. This latter analysis demonstrates that the significant decline detected was specific to known hotspot motifs; declines of similar degree were rarely observed in non-hotspot motifs (**Supplemental File 1**).

### A codon substitution model for phylogenies undergoing somatic hypermutation

All three bNAb lineages showed a significant improvement in likelihood under the symmetric WR<u>C</u>/<u>G</u>YW HLP16 model compared to the GY94 model. The maximum likelihood values of *h* for the three data sets were *ĥ*^*WR<u>C</u>*^ = *ĥ*^*<u>G</u>YM*^ = 1.91, 1.82, and 2.05, for CH103, VRC26, and VRC01, respectively. In each case the simpler GY94 model (all *h=0*) could be rejected using the likelihood ratio test (*p* < 0.0001 for all three lineages). These results are summarized in **Table 2**. These *ĥ* values represent up to a three-fold increase in the relative rate of change at hotspot locations (depending on the values of *b*_*ij*_).

**Table 2:** Maximum likelihood estimates of *h* and likelihood ratio tests

| Lineage | $\hat{h}^{WRC/GYW}$ | Log-likelihood | | 2*LR | p value |
| --- | --- | --- | --- | --- | --- |
| | | $\hat{h}^{WRC/GYW}=\text{mle}$ | $\hat{h}^{WRC/GYW}=0$ | | |
| CH103 | 1.91 (1.5, 2.4) | -14927 | -15031.5 | 209 | 0 |
| VRC26 | 1.82 (1.6, 2.1) | -37632.5 | -37913.8 | 562.6 | 0 |
| VRC01 | 2.05 (1.8, 2.3) | -44037.7 | -44339.3 | 603.2 | 0 |

Significance was determined using the likelihood ratio test under a chi-squared distribution with one degree of freedom. 90% confidence intervals for $\hat{h}$ are shown in parentheses in the second column. All lineages showed a p value $< 1 \times 10^{-45}$ .

The mean field approximation used in this model did not dramatically affect parameter estimation when applied to data sets simulated under a fully context dependent model, at least for the parameter space of the three empirical bNAb lineages (**Table 3**). Mean *ĥ* values from simulations in which 0 ≤ *h^WR<u>C/G</u>YW^* ≤ 2 were close to their true *h* values and exhibited low absolute bias and variability (maximum-0.17 and 0.11, respectively, when *h* =2). Of these simulated data sets, 6.1% incorrectly rejected the correct parameter value (i.e. they estimated a *ĥ* significantly different from the true value of *h* used in the simulations). This is close to the theoretical expectation under α = 0.05. Further, none of the datasets simulated with *h* > 0 failed to reject the null hypothesis that *h* = 0, demonstrating good statistical power. Bias generally increased if *h* was raised beyond that observed in the empirical bNAb linages. Performance was worse when *h* = 4, which resulted in a mean type 2 error of 0.42 and a mean bias of -0.59. This behaviour is as expected because, as *h* increases, the mean field approximation will become less accurate. We found that using empirical codon frequencies decreased the performance of *h* estimation; using empirical frequencies resulted in higher bias and type 2 error rates than using ML frequencies (**Supplemental File 3**). Discussion of why empirical codon frequencies are unlikely to be suitable for long-lived B-cell lineage phylogenies is provided in the Methods section.

**Table 3:**
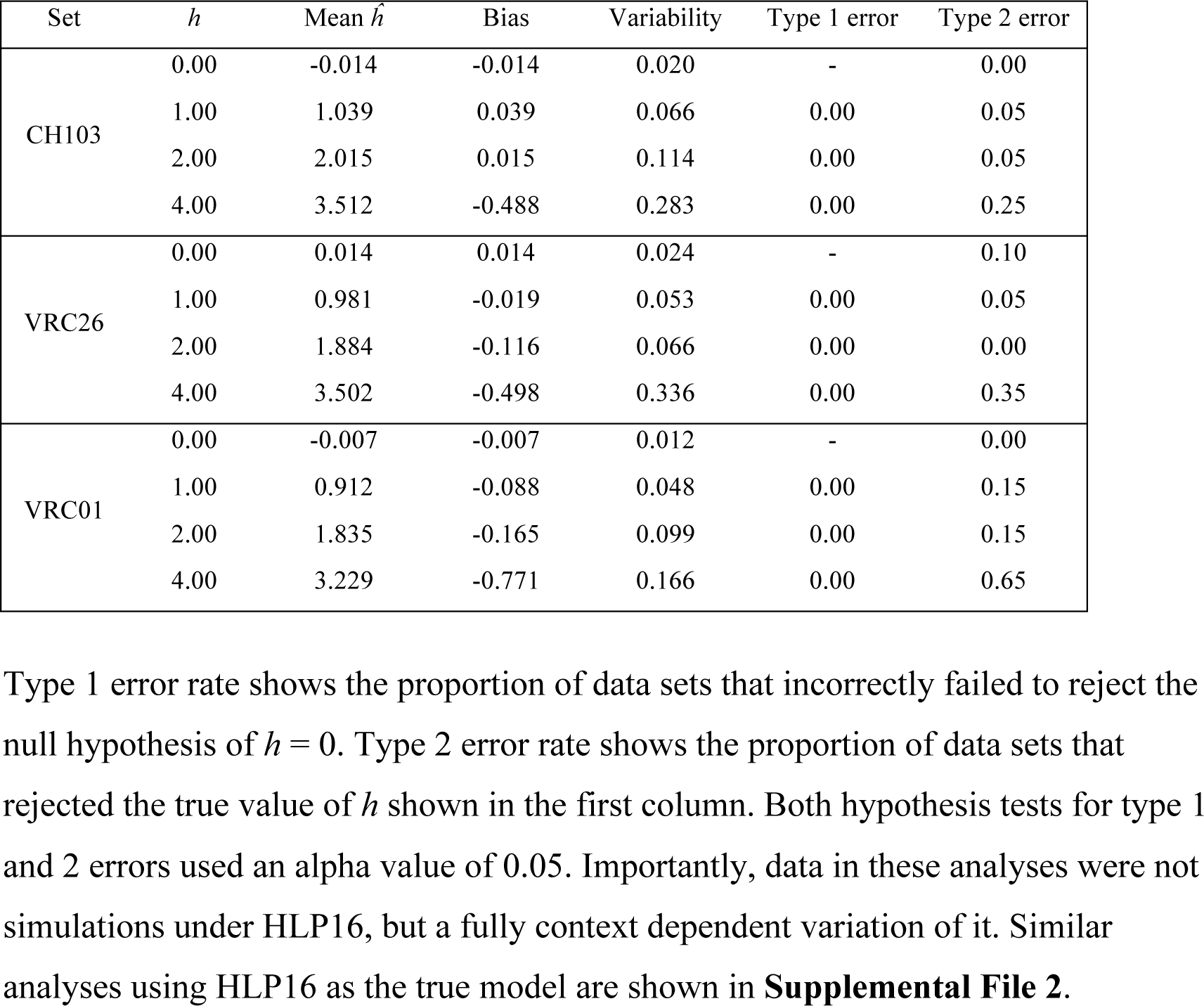
HLP16 performance under fully-context dependent simulations for symmetric WR<u>C</u>/<u>G</u>YW hotspots

**Table 4:** Hotspot model selection

| Constraint/optimization of each $h^a$ | | | | | | | $p$ values from $LR$ tests | | |
| --- | --- | --- | --- | --- | --- | --- | --- | --- | --- |
| Model name | $h^{WRC}$ | $h^{GYW}$ | $h^{WA}$ | $h^{TW}$ | $h^{SYC}$ | $h^{GRS}$ | CH103 | VRC26 | VRC01 |
| Symmetric WRC/GYW* | ML | $h^{WRC}$ | 0 | 0 | 0 | 0 | 2.3E-15 | 7.8E-05 | 3.8E-03 |
| Asymmetric WRC/GYW | ML | ML | 0 | 0 | 0 | 0 |  |  |  |
| Symmetric WA/TW* | 0 | 0 | ML | $h^{WA}$ | 0 | 0 | 0 | 0 | 0 |
| Asymmetric WA/TW | 0 | 0 | ML | ML | 0 | 0 |  |  |  |
| Symmetric SYC/GRS* | 0 | 0 | 0 | 0 | ML | $h^{SYC}$ | 0.65 | 2.2E-13 | 4.2E-03 |
| Asymmetric SYC/GRS | 0 | 0 | 0 | 0 | ML | ML |  |  |  |
| Uniform hotspots* | ML | $h^{WRC}$ | $h^{WRC}$ | $h^{WRC}$ | 0 | 0 | 6.7E-16 | 0 | 0 |
| Hierarchical hotspots | ML | $h^{WRC}$ | ML | $h^{WA}$ | 0 | 0 | | | |
| SCAH* | ML | ML | ML | ML | ML | $h^{SYC}$ | 0.65 | 1.1E-06 | 1.1E-03 |
| FCH | ML | ML | ML | ML | ML | ML |  |  |  |

Models of hotspot hierarchy (degree of mutability) and symmetry, specified by placing constraints on how different values of $h$ are optimized. Columns 2-7 show how the parameter $h^a$ is obtained for a particular model. A value of “0” indicates that $h$ is fixed at zero, “ML” indicates that a parameter is optimised by maximum likelihood, and “ $h^a$ ” indicates that $h$ parameter is equal to another value of $h$ . For instance, in “Symmetric WRC/GYW,” $h^{GYW}$ is equal to its reverse complement $h^{WRC}$ , which is ML optimised. However, in “Asymmetric WRC/GYW,” both are ML optimised. Note that each model marked with an asterisk \* is nested with the model immediately below it by one free parameter, allowing hypothesis testing using a likelihood ratio test. Rows 8-10 show $p$ values obtained from likelihood ratio tests of each of these nested hotspot models for the bNAb lineage specified in each column. SCAH = symmetric coldspots, asymmetric hotspots; FCH = free coldspots and hotspots. Parameters, log likelihood, and AIC of each fit are shown in **Supplemental File 5**.

Within the parameter space of the empirical data sets (0 ≤ *h^WR<u>C/G</u>YW^* ≤ 2), there was no substantial difference in estimation of other model parameters compared to the standard GY94 model, except for the tree length parameter in some simulations **(Figure 2**, **Supplemental File 4**). However, when this *h* is large (4, in these simulations), the GY94 model substantially underestimates tree length in each of the simulated lineages. In contrast, the HLP16 model, while not completely eliminating this effect, substantially reduced it. In simulations based on the long-lived VRC01 lineage in which this *h* = 4, the GY94 model overestimated the ω parameter; this bias was not obvious in simulations based on the VRC26 and CH103 lineages that were sampled for a shorter duration. The HLP16 model was generally able to infer ω accurately under all values of *h*.

### Hotspot model selection

All hotspot motif models tested gave a significantly higher likelihood than the standard GY94 model when applied to the CH103, VRC26, and VRC01 B-cell lineages. Likelihoods were considerably higher for asymmetric models. Using a LRT, the asymmetric WR<u>C</u>/<u>G</u>YW model significantly rejected the corresponding nested symmetric model (p = 2.3x10^−15^, 7.8x10^−5^, and 3.8x10^−3^, for lineages CH103, VRC26, and VRC01, respectively). Similarly the asymmetric W<u>A</u>/<u>T</u>W model rejected its symmetric counterpart (p < 1x10^−45^ for all three lineages). Allowing different hotspot motifs to have different *h* values also resulted in significantly higher likelihoods than using a uniform value of *h* for all hospots (p < 1x10^−15^ for all three lineages). Interestingly, VRC26 and VRC01 showed a significantly higher likelihood under asymmetric SY<u>C</u>/<u>G</u>RS coldspot motifs (p = 2.2x10^−13^ and 4.2x10^−3^), but CH103 did not (p=0.65). This difference was also reflected in the best-fit (lowest AIC) model for each lineage. For VRC26 and VRC01the best-fit model was the “Free coldspots and hotspots” model, in which all motifs and their reverse complements are given separate *h* values. However, for CH103 the best-fit model was the “Symmetric coldspots, asymmetric hotspots” model, in which each hotspot and its reverse complement are given separate *h* values, but coldspots remain symmetric.

In the randomization analysis, we found that WR<u>C</u>/<u>G</u>YW motifs exhibited a larger value of *h*, and a higher likelihood, than any other trimer motif analysed. Further, SY<u>C</u>/<u>G</u>RS motifs resulted in a *h* values that was lower than 140 of the 143 other trimer motifs tested. W<u>A</u>/<u>T</u>W motifs showed a higher *h* value than 22 out of the 23 other dimer motifs analysed (only R<u>C</u>/<u>G</u>Y motifs showed a higher *h*). These results are shown in **Supplemental File 6**.

### Ancestral state reconstruction

In fully context dependent simulations, we also found that the HLP16 model provided an accuracy of ancestral state reconstructions that was similar to the GY94 model where *h* < 4, and that HLP16 substantially improved accuracy at *h* = 4 (**Supplemental File 7**). Sequence reconstructions under the two models were fairly similar for internal nodes near the root and the tree tips, but showed improvement under the HLP16 model especially for internal nodes in the basal third of the phylogeny. Typically, we would expect the uncertainty in ancestral state reconstruction to increase as we move from the tree tips towards the root; however, B-cell lineages are unusual in that the root sequence is also known as it corresponds to the germline sequence.

While true ancestral sequences are not available for the three empirical bNAb lineages, we did observe differences between ancestral sequences reconstructed using the HLP16 and GY94 models. For each lineage, we compared the two models by calculating the mean number of amino acid differences between the predicted ancestral sequences at all internal nodes of each tree. Performing this ancestral state reconstruction on each of the three bNAb lineages showed a mean of 0.63, 1.15, and 0.95 amino acid sequence difference across all internal nodes, with a maximum difference of 9, 10, and 15 amino acid differences in a single node for CH103, VC26, and VRC01, respectively. Differences somewhat more concentrated in the basal third of the phylogeny, consistent with the simulation results above (**Supplemental File 8**).

## Discussion

Molecular phylogenetics has already been used in a variety of applications in the study BCR genetic diversity and the molecular evolution of B cell lineages (Kepler 2013; Sok et al. 2013; Kepler et al. 2014). However, the process of somatic hypermutation is known to occur in ways that violate fundamental assumptions of most phylogenetic substitution models. Here, we demonstrate that failing to account for this has tangible effects on phylogenetic inference and ancestral state reconstruction from sets of sequences from long-lived bNAb lineages. We develop and implement a new codon substitution model (HLP16) that, whilst only an approximation, is capable of mitigating these effects.

Perhaps the most salient difference between standard substitutions models and the biology of somatic hypermutation is the context dependency of mutation in BCRs. This biased mutation process at hotspot motifs, for which a variety of empirical models have been developed to characterise the process at di, tri, penta, and heptamer levels (Smith et al. 1996; Yaari et al. 2013; Elhanati et al. 2015), has long been known to give false signature of selection in BCRs (Dunn-Walters and Spencer 1998). This effect was observed in some of our simulations (**Figure 2**, **Supplemental File 4**), as a failure to account for the increased rate of substitution at hotspot motifs led to overestimation of the ω (dn/ds) parameter. However, these simulations used an *h* value of 4, which was outside of the range of what we observed for empirical bNAb lineages.

Some approaches have been developed to study the substitution process in BCR data in the context of biased mutation. Some of these are non-phylogenetic in nature (e.g. Hershberg et al. 2008; Yaari et al. 2012) and focus on the expected number of germline to tip replacement mutations in comparison to a null model. Kepler et al (2014) developed a non-linear regression model approach that, combined with an empirical model of mutation rate at each site, allowed the authors to test for the effects of selection and mutation on BCR genetic diversity. The substitution model detailed in McCoy et al (2015) is more similar to the model introduced in our study, but accounts for biased mutation by comparing values of ω inferred from a given data set to those inferred from out-of-frame rearrangements.

Other approaches have been taken to study the effect of context dependent mutation in phylogenetic substitution models. Many have focused on modelling the substantially increased mutability of CpG motifs (Hwang and Green 2004; Lunter and Hein 2004). These approaches are attempts to account for the full context dependency of CpG hypermutation, and require significantly more complex models. In the case of somatic hypermutation in BCRs, the increased mutability of BCR hotspot mutations (~3 fold) is not as great as CpG motifs (~18 fold; Lunter and Hein 2004), so a simpler, approximate approach is still effective (**Table 3**). The mean-field approximation has also been used previously, but in a reversible codon model, to take into account di- and trinucleotide substitutions (Whelan and Goldman 2004).

The HLP16 codon substitution model detailed here is a relatively straightforward modification of the widely used M0 submodel of the GY94 model. Although the HLP16 model is slower to compute than the simpler, reversible model on which it is based, we have found that it is usable, and certainly statistically preferable, to the GY94 model when applied to any BCR data set whose diversity may have been shaped by somatic hypermutation. Further, the HLP16 model does not rely on an empirical model to incorporate the effect of biased mutation, but instead attempts to explicitly model the context-dependent mutational process by estimating the parameter *h* directly from the data. We note, however, that the HLP16 model is a mean-field approximation and does not capture the full context of motif driven evolution. Therefore we do not expect it to fully disentangle interactions between selection and biased mutation, and estimated values of ω should be interpreted carefully. In addition to correcting biases in parameter estimation, simulation analyses reveal that the HLP16 model produces different, and more accurate, ancestral state reconstructions than the standard GY94 model. Importantly, empirical analyses on bNAb lineages performed here were using tree topologies that were optimal under GY94, rather than HLP16, for computational tractability. This is expected to make the estimation of each *h* conservative in these analyses, but it is not clear how the optimal topology of the HLP16 model will differ from that under GY94.

Our model selection results suggest that different hotspot motifs have highly variable effects on sequence evolution in B-cell lineages. It is generally thought that increased mutation in WRC/GYW motifs (or the tetramer motifs WRCY/RGYW) reflect the action of AID targeting, while in WA/TW motifs it is the result of error-prone polymerase repair (Teng and Papavasiliou 2007). Consistent with these separate mechanisms, WRC/GYW motifs have generally been found to be strand symmetric, but WA/TW motifs are strand-biased, with WA mutating at a higher rate than TW (Bransteitter et al. 2004; Spencer and Dunn-Walters 2005; Teng and Papavasiliou 2007). It is interesting, then, that in all three lineages tested here show a significantly better fit for asymmetric vs. symmetric WRC/GYW (**Table 4; Supplemental File 5**). However, our results do not necessarily conflict with previous findings on the targeted nature of SHM. If strand bias were a feature of AID targeting, it would be expected to be consistent between lineages. However, the asymmetric WRC/GYW model did not show a consistent polarity, with CH103 and VRC01 having *h^GYW^* > *h^WRC^*, and VRC26 showing the opposite pattern (**Supplemental File 5**). By contrast, the asymmetric WA/TW model also showed a higher value of *h^WA^* than *h^TW^*, consistent with the existing literature. One can imagine a number of complex factors that may lead to increased likelihood under the asymmetric WRC/GYW model even under a strand symmetric targeting of AID, and these tests do not distinguish between them. SYC/GRS coldspot motifs also did not show a consistent strand polarity between the lineages, and in CH103 did not show evidence of asymmetry at all, consistent with the notion that SYC/GRS motifs are also the result of AID targeting (Bransteitter et al. 2004).

Another common assumption in phylogenetic analysis is that the codons or nucleotides sampled for analysis are at their equilibrium frequencies. Because our hotspot model has asymmetric relative rates between codons, which are a function of *h*, codon frequencies may change through time within a B-cell lineage when *h* is significantly above zero. This is a consequence of the decline in the number of hotspots through time (**Figure 1**). We dealt with this problem by estimating equilibrium frequencies by maximum likelihood within the model. This provided an improvement, both in maximum likelihood and in parameter estimation, over using empirical codon frequencies. However, it is not yet clear if this is the most efficient or the most effective way of dealing with the problem sequences that have not converged to their equilibrium distribution. While ML optimization finds the best fitting codon frequency values (under the CF3x4 model), in reality codon frequencies may change over the course of the phylogeny, and a model that accounts for that would likely be more appropriate. However, effective modelling of the numerous factors that affect codon frequency change in BCR lineages will be complex and we leave that problem for future analyses.

This decay of hotspot motifs in bNAb lineages may have important implications for our understanding of host-virus coevolution. More specifically, the loss of hotspot motifs may lead to a decrease in sequence mutability, and therefore a decline in overall rate of evolution over time for a given lineage (Sheng et al. 2016). This hypothesis has several interesting implications. If the slowdown in mutation rate over time, arising from hotspot decay, is an intrinsic property of activated B cell lineages, then BCR sequence divergence from a germline ancestor (and thus affinity maturation) may be intrinsically constrained. Consequently, while BCR lineages may be able to rapidly evolve binding affinity and co-evolve with pathogens for an initial period after activation, over longer periods of time the ratio of the rate of BCR sequence change to pathogen sequence change may decline. We hypothesise that in extreme cases the rates of BCR evolution within a lineage may eventually fail to keep up with the rapid evolution of chronically infecting viruses, such as HIV-1, due to the exhaustion of available BCR hotspot motifs. The notion that biased mutation will lead to decreased mutability and evolutionary rate was explored recently by Sheng et al (2016). They concluded that the observed mutation rate decreases in bNAb lineages was most likely due to a shift from positive to purifying selection, although the loss of hotspot motifs may also play a role and the issue is not yet fully resolved.

We have implemented this model in the software package IgPhyML, a modified version of codonPhyML (Gil et al. 2013). This program can perform all of the substitution model analyses performed here. Source code is available at: https://github.com/kbhoehn/IgPhyML.

## Acknowledgements

This work was funded by the European Research Council under the European Union’s Seventh Framework Programme (FP7/2007-2013)/ERC grant agreement no. 614725-PATHPHYLODYN. KBH was also supported by a Marshall scholarship.

